# Temporal requirements of MAPK effectors reflect signalling microenvironment heterogeneity during *Mesp1*+ cardiac progenitor emergence and migration

**DOI:** 10.1101/2025.08.13.670016

**Authors:** Nitya Nandkishore, Sinem Eylem Inal, Adeline Ghata, Fabienne Lescroart

## Abstract

During mouse gastrulation, cardiac progenitors arise within dynamic morphogenetic landscapes, yet the mechanisms by which the progenitors integrate this information and respond to it remain unclear. We profiled candidate signalling ligands in wild-type and *Mesp1* mutant embryos, which display defective progenitor migration. In Mesp1⁺ progenitors, *Fgf* and *Wnt* ligand expression changes over gastrulation, coinciding with the emergence of first and second heart fields (FHF, SHF). This temporal pattern is perturbed in *Mesp1* mutants. Signal inhibition studies in embryos, cultured *ex utero,* reveal a role for intracellular signalling effectors in mediating the proper migration of the cardiac progenitors. We demonstrate that the different branches of the MAPK signalling effectors (p38, ERK, JNK) are required in distinct temporal windows for proper migration. Our findings link the heterogeneous signalling environment of the cardiac fields to stage-specific MAPK requirements, offering insight into how progenitors integrate complex environmental inputs to coordinate fate specification and migration.

## Introduction

Organogenesis requires the precise specification and coordinated deployment of progenitor cells, and perturbations in these processes can lead to congenital malformations. Congenital heart defects (CHDs) are a prominent example, affecting approximately 1 in about 100 live births (Bajolle et al., 2009; Kloesel et al., 2016). Elucidating the earliest molecular and cellular events underlying the specification and deployment of cardiac progenitors is essential for advancing our understanding of CHD pathogenesis.

### Heart development originates from a heterogenous pool of *Mesp1*− expressing progenitors during gastrulation

The heart is formed by progenitor cells that arise during gastrulation from the primitive streak (PS) and are marked by the expression of the key transcription factor MESP1 (Lescroart et al., 2014; Saga, 2000; Saga et al., 1999). *Mesp1*− expressing cells leave the PS progressively during the time course of gastrulation and exhibit lineage restriction toward specific cardiac cell types and compartments (Devine et al., 2014; Ivanovitch et al., 2021; Lescroart et al., 2014). The earliest emerging progenitors, corresponding to the first heart field (FHF) or the partially overlapping juxta-cardiac field (JCF), preferentially contribute to the left ventricular (LV) myocardium (Lescroart et al. 2018; Abukar et al., 2023; Ivanovitch et al., 2021; Kelly, 2023; Lescroart et al., 2014; Tyser et al. 2021; Zhang et al. 2021). Progenitors that emerge later correspond to the second heart field (SHF), which can be broadly divided into two populations. Anterior SHF (aSHF) progenitors, arising from the distal PS, contribute to the right ventricular (RV) myocardium and arterial pole (OFT) of the heart (Abukar et al., 2025; Ivanovitch et al., 2021; Kelly, 2023; Lescroart et al., 2014), while posterior SHF (pSHF) progenitors arising from the proximal PS, contribute to the atria (LA/RA) and venous pole of the heart (Fig. 1a).

**Figure 1:**
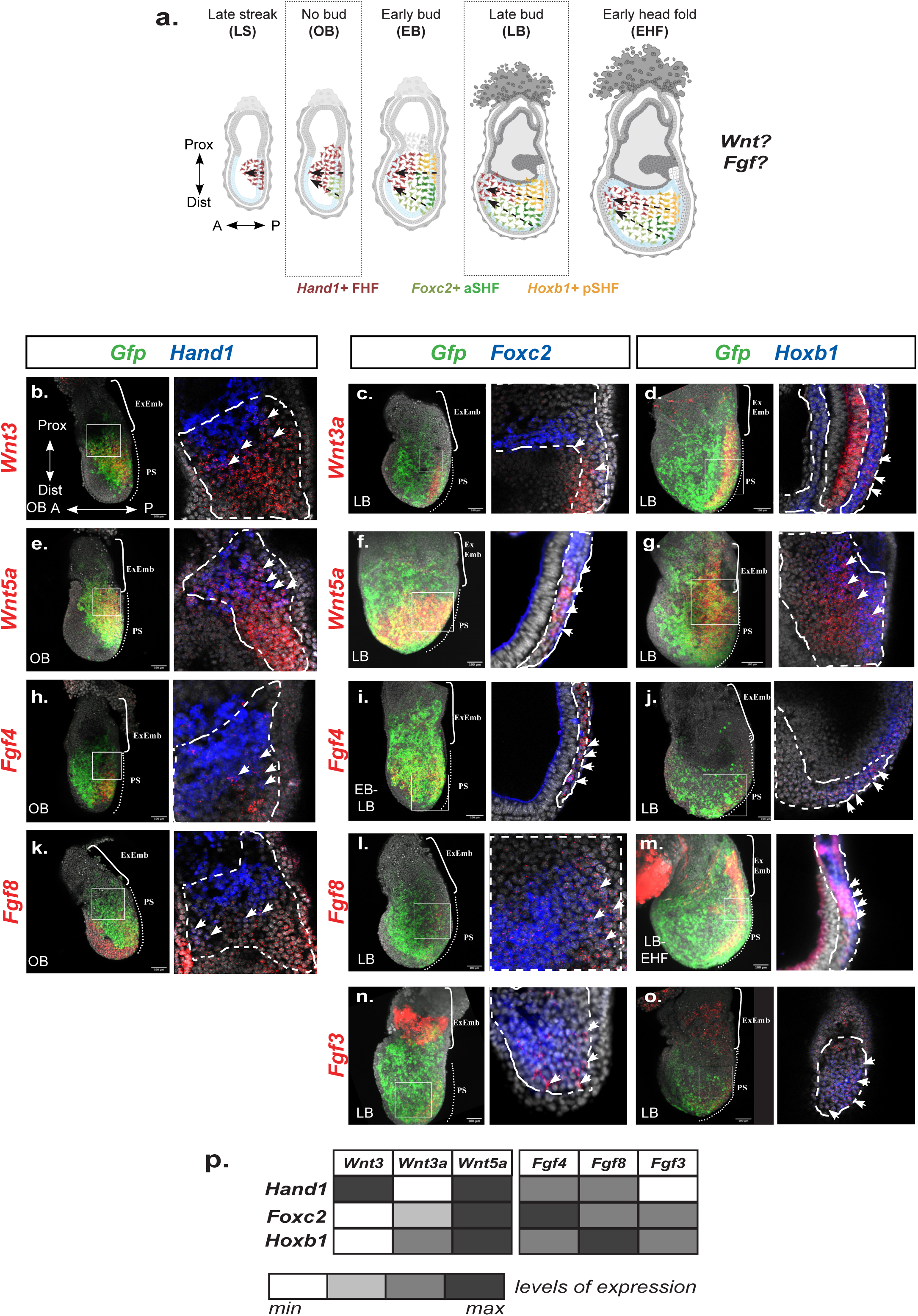
*Fgf* and *Wnt* expression in *Mesp1+* progenitors evolves during gastrulation. **a.** Scheme of the migration of the different subpopulations of *Mesp1+* cardiac progenitors (*Hand1+* FHF in red, *Foxc2+* aSHF in green and *Hoxb1+* pSHF in yellow) during gastrulation at the late streak (LS), No Bud (OB), Early Bud (EB, Late Bud (LB) and early head forld stages (EHF). **b-o.** Representative maximum intensity projection images of *Mesp1-rtTA/tetO-H2B-GFP* embryos at the OB (**b**, **e**, **h**, **k**) or LB stage (**c-d**, **f-g**, **i-j**, **l-m**, **n-o**) after RNAscope experiment with *Gfp*, *Wnt3* (**b**), *Wnt3a* (**c-d**), *Wnt5a* (**e-g**), *Fgf4* (**h-j**), *Fgf8* (**k-m**) and *Fgf3* (**n-o**) with respect to *Hand1*, *Foxc2* or *Hoxb1* expression. *Gfp* marks *Mesp1+* progenitors. Images on the left are maximum projections describing the pattern of expression of the signalling ligand (in red) with respect to the *Mesp1-GFP* expressing cells (in green), while images on the right are single optical sections magnified from the region indicated with a white box, describing the pattern of expression of the ligand with respect to the marker of the different subpopulations (in blue). White dashed lines in the single optical sections indicate the region of *Mesp1-GFP* expressing cells. White arrowheads indicate representative individual cells that express both *Hand1, Foxc2 or Hoxb1* and the ligand in a single optical section. **p.** Summary table to describe the qualitative expression of the various intercellular signalling ligands in the different cardiac subpopulations. Levels of expression (min-minimum, max-maximum) are represented with nuances of grey. A, anterior; P, posterior; Prox, proximal; Dist, distal; ExEmb, extra-embryonic region (square bracket); PS, Primitive streak (dotted line). Scale bars: 100μm. Each embryo is a representative of at least 3 embryos at the defined stage.

### Cardiac progenitors are deployed through different migratory patterns

After their emergence from different levels of the PS, *Mesp1*− expressing cells migrate laterally and anteriorly toward the anterior pole of the embryo (Fig. 1a) where they will form the first differentiated structure, the cardiac crescent (Buckingham et al., 2005; Kelly, 2016). Recent advances in live imaging of mouse embryos have revealed that the pattern of migration is actually different for the distinct subpopulations of cardiac progenitors (Dominguez et al., 2023; Abukar et al., 2025). Notably, there is a correlation between their route of migration and their ultimate cardiac fate.

### Cell migration is directed by MESP1 during gastrulation

Little is known about the mechanisms governing early cardiac progenitor migration. A role for MESP1 has been established, as *Mesp1* mutant embryos exhibit migration defects (Dominguez et al., 2023; Saga et al., 1999), with progenitors failing to migrate from the PS (Saga et al., 1999). *In vitro* studies have further demonstrated that MESP1 controls the speed, polarity and directionality of cardiac progenitor migration (Chiapparo et al., 2016). Moreover, components of the WNT and FGF signalling pathways, known to guide cardiac cell migration (Münsterberg and Yue, 2008; Song et al., 2014; Sweetman et al., 2008; Yang et al., 2002; Yue et al., 2008) have been identified as downstream targets of MESP1. These targets are also dysregulated in *Mesp1* mutant embryos (Lescroart et al., 2018; Lin et al., 2022). However, how different subpopulations of *Mesp1*+ cardiac progenitors adopt distinct migratory routes and fates remains poorly understood. While single cell transcriptomics have identified intercellular signalling ligands and transcription factors that are differentially expressed among the various cardiac subpopulations (Krup et al., 2023; Lescroart et al., 2018), the role of downstream intracellular signalling effectors in integrating and amplifying these cues to produce a coherent cellular response has not been thoroughly investigated.

In this study, we investigated the expression of few *Fgf* and *Wnt* genes enriched in cardiac progenitors and whether intracellular signalling effectors are differentially required for the development of distinct cardiac progenitor subpopulations. We first assessed whether the signalling environment differs at the time and location of emergence of each subpopulation. We then examined how this environment is altered in *Mesp1* mutant embryos. Finally, we tested whether the requirement for intracellular signalling effectors varies spatially and temporally at the onset of cardiac progenitor specification.

## Results

### *Fgf* and *Wnt* expression in Mesp1+ progenitors evolves during gastrulation

Transcripts encoding for the ligands of the WNT and FGF pathways are enriched in *Mesp1*− expressing progenitors, and are downstream targets of MESP1 (Lescroart et al., 2018, 2014; Lin et al., 2022). While *Wnt* and *Fgf* ligand expression in the early embryo has been previously described (Arnold and Robertson, 2009; Chen et al., 2019; Du et al., 2010; Kimura et al., 2001; Kinder et al., 1999; Niswander and Martin, 1992; Nowotschin et al., 2012; Sun et al., 1999; Tremblay et al., 2001; Yamaguchi et al., 1999), we sought to examine their expression in the context of the distinct *Mesp1*+ cardiac progenitor subpopulations. We hypothesized that these subpopulations emerge within distinct signalling microenvironments.

We used a tetracycline-inducible *Mesp1* reporter transgenic mouse line, *Mesp1-rtTA/tetO-H2B-GFP* (Lescroart et al., 2014), and induced GFP reporter expression at the onset of gastrulation (E6.25). Embryos were collected at either early (no bud, OB at around E6.75) or late (late bud, LB at around E7.25) gastrulation stages (Fig. 1a). We analysed the expression of *Wnt* and *Fgf* ligand encoding genes, that are enriched in *Mesp1+* progenitors (Lescroart et al., 2014), using RNAscope in combination with markers of distinct cardiac progenitor subpopulations: *Hand1,* for the prospective FHF/JCF (Lescroart et al., 2018; Tyser et al., 2021; Zhang et al., 2021) (Supplementary Fig. 1a-b); *Foxc2*, for the aSHF and distal/anterior PS progenitors (Kaestner et al., 1996; Lescroart et al., 2018) (Supplementary Fig. 1c), and *Hoxb1*, for the pSHF and proximal/posterior PS progenitors (Bertrand et al., 2011; Lescroart et al., 2018) (Supplementary Fig. 1d). RNAscope revealed that *Wnt3* is expressed in emerging progenitors during early gastrulation (OB stage), with few cells co-expressing *Hand1*, but its expression is downregulated at later stages (EB stage) (Fig. 1b, Supplementary Fig. 1e, h). In contrast, *Wnt3a* expression is detected after the onset of *Wnt3* expression (Fig. 1c-d) and becomes restricted to the PS at the LB stage (Fig. 1c-d, Supplementary. Fig 1i). *Wnt3a* is moderately co-expressed with *Foxc2* and more strongly with *Hoxb1*. *Wnt5a* is expressed in emerging *Mesp1*+ cardiac progenitors at both early (OB) and later (LB) gastrulation stages, in co-expression with *Hand1*, *Foxc2* and *Hoxb1* (Fig. 1e-g, Supplementary Fig. 1j). These findings indicate that spatiotemporally different *Mesp1*+ subpopulations are exposed to distinct WNT ligands during gastrulation. *Wnt3* is specific to the FHF/JCF progenitors while *Wnt3a* is more enriched in anterior and posterior SHF progenitors.

After demonstrating the dynamic expression of *Wnt* ligands, we next explored the spatial and temporal expression of *Fgf* ligand encoding genes in Mesp1+ subpopulations. *Fgf4* is expressed by progenitors emerging along the PS early in gastrulation (OB stage), with co-expression in *Hand1*-positive cells (Fig. 1h-j, Supplementary Fig 1k), and appears enriched in progenitors from the anterior PS. This is particularly evident at later stages (LB stage), when its expression is higher in progenitors emerging from *Foxc2*-positive aSHF progenitors compared to *Hoxb1*-positive pSHF progenitors (Fig. 1i-j). *Fgf8* is broadly expressed in all germ layers at early gastrulation stages (Fig. 1k, Supplementary Fig. 1l), but becomes downregulated and restricted to the PS later (Fig. 1k-m), consistent with its role in cell migration (Andre et al., 2015; Hardy et al., 2008; Yamaguchi et al., 1999; Ye et al., 2013). *Fgf8* is co-expressed with *Hand1*, *Foxc2* and Hoxb1, with the highest expression in the *Hoxb1*-positive subpopulation. *Fgf3* is absent in the embryonic region at the earliest stages of gastrulation (Supplementary Fig. 1f) but is expressed by progenitors emerging later during gastrulation, in co-expression with *Foxc2* and *Hoxb1* (Fig. 1n-o, Supplementary Fig. 1m). These findings further support the idea that genes encoding FGF ligands are heterogeneously expressed in Mesp1+ subpopulations, with *Fgf4* enriched in *Foxc2*-positive cells, *Fgf8* in *Hoxb1*-positive cells, and *Fgf3* absent from *Hand1*-positive progenitors (Fig. 1p).

Taken together, and in the context of the cardiac progenitor subpopulation identity (Ivanovitch et al., 2021; Kelly, 2023; Lescroart et al., 2014), these results suggest that early-emerging *Hand1+/Mesp1*+ cardiac progenitors (FHF/JCF/LV) arise in an environment enriched in *Wnt3, Wnt5a, Fgf4* and *Fgf8*. Later-emerging cardiac progenitors appear to experience a signalling microenvironment comprising *Wnt5a* and *Fgf3* and in addition, *Foxc2+* progenitors from the distal PS (aSHF/RV/OFT) are exposed to high *Fgf4*, moderate *Fgf8* and lower *Wnt3a* mRNA levels, while *Hoxb1+* progenitors from the proximal PS (pSHF/LA/RA) are exposed to high *Fgf8*, moderate *Fgf4,* and *Wnt3a* mRNA levels (Fig. 1p).

### *Fgf* and *Wnt* expression is altered in *Mesp1* mutants

Given the heterogeneity of signalling from the microenvironment during gastrulation, we sought to determine whether this environment is altered in *Mesp1* mutants, which display defects in cardiac progenitor migration. We crossed the *Mesp1-Cre* mutant allele into the background of the same *Mesp1* reporter mouse line used previously (Fig. 2a) and analysed the expression of *Fgf* and *Wnt* ligands by RNAscope. Mutant embryos were compared to litter- and stage-matched heterozygous controls. The mutant embryos could also be distinguished by the accumulation of *Mesp1-*GFP cells at the posterior pole, as expected in the *Mesp1* mutants (Dominguez et al., 2023; Saga et al., 1999). Consistent with previous single-cell RNA sequencing data (Lescroart et al., 2018), *Fgf8* expression was found to be upregulated in *Mesp1* mutants (Fig. 2b-c). In contrast, *Fgf3* expression is lost in the embryonic region (Fig. 2d-e). Similarly, we observed a loss of *Wnt3* expression in *Mesp1* mutant embryos (Fig. 2f-g). *Wnt5a* expression was downregulated and became restricted to the *Mesp1-GFP*+ cells (Fig. 2h-i). By contrast, the initiation of *Wnt3a* expression appeared unaffected (Fig. 2j-k). These results indicate that the signalling microenvironment of cardiac progenitors is significantly altered in *Mesp1* mutants compared to wild-type embryos at equivalent stages. Specifically, we observed increased *Fgf8*, and loss of *Fgf3* and *Wnt3* expression (Fig. 2l).

**Figure 2:**
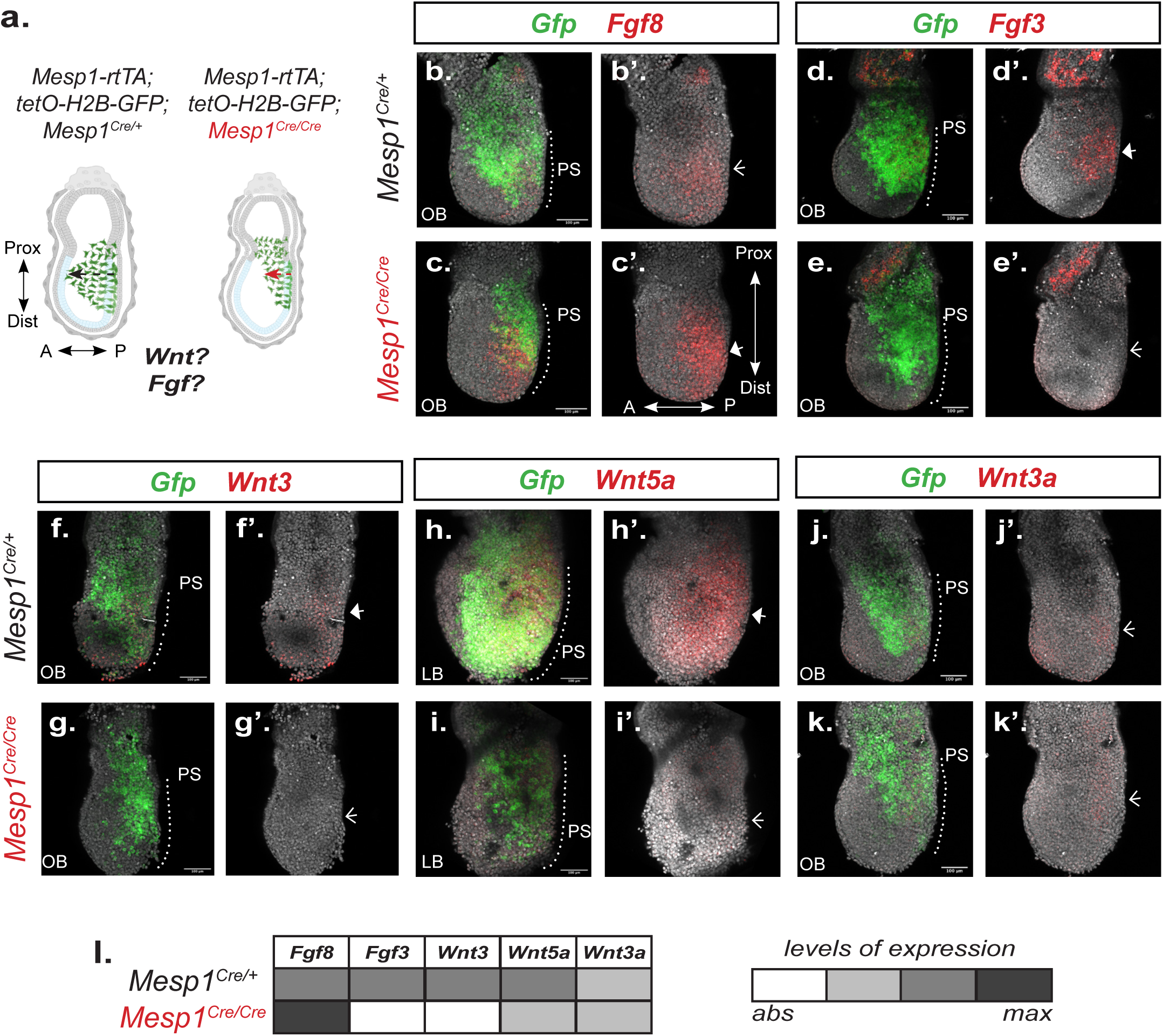
*Fgf* and *Wnt* expression is altered in *Mesp1* mutants. **a.** Experimental scheme of the experiment. *Mesp1* lineage is tracked in *Mesp1-rtTA/tetO-H2B-GFP; Mesp1^Cre/Cre^* embryos after doxycycline induction with the stages of collection clearly defined. *Mesp1^Cre/Cre^* mutants show migration defects (right). **b-k.** Representative maximum intensity projection images of embryos after RNAscope for *Fgf8* (**b-c**), *Fgf3* (**d-e**), *Wnt3* (**f-g**), *Wnt5a* (**h-i**) and *Wnt3a* (**j-k**) with respect to the *Mesp1+ (Gfp)* expressing cells during gastrulation in *Mesp1* heterozygous (Cre/+) (**b, d, f, h, j**) and *Mesp1* mutant (Cre/Cre) (**c, e, g, i, k**) embryos at no bud (OB) or late bud (LB) stages of gastrulation (E7.25). Scale bars: 100μm. A, anterior; P, posterior; Prox, proximal; Dist, distal; PS, Primitive streak (dotted line). All embryos are represented in the same orientation. **b’-k’** represent the same images with only the expression pattern of the *Fgf* or *Wnt* ligand. Arrowheads indicate domains of ligand gene expression with solid arrowheads indicating domains of higher gene expression. Each embryo is a representative of at least 3 embryos. **c.** Summary table to describe the qualitative levels of expression (abs-absent, max-maximum) of the *Fgf* and *Wnt* ligands in *Mesp1* heterozygous (Cre/+) and *Mesp1* mutant (Cre/Cre) embryos.

### Mitogen-activated protein kinases (MAPKs) mediate signalling in *Mesp1+* cardiac progenitors

We have shown that at the time of emergence of the distinct cardiac progenitor populations, the signalling microenvironment is heterogenous, with subtle differences in ligand encoding gene expression across subpopulations. Despite the variety in ligands within each signalling pathway, the diversity of receptors they activate is limited (Qin et al., 2024; Teven et al., 2014). However, these limited receptors can activate a range of downstream intracellular signalling effectors to elicit specific cellular responses. These effectors not only amplify the intracellular signal but may also function as nodes integrating multiple pathways to generate context specific outcomes (Dailey et al., 2005; Qin et al., 2024). With this in mind, we focused on intracellular signalling effectors potentially mediating differential responses among cardiac progenitor subpopulations. We selected the mitogen-activated protein kinase (MAPK) signalling pathway, given its established role in cell migration and proliferation (Huang et al., 2004; Zhang and Liu, 2002) – two critical processes during gastrulation. MAPK signalling is also known to act downstream of several intercellular signalling ligands expressed during gastrulation; including – but not limited to – FGF and WNT ligands (Bikkavilli and Malbon, 2009; Clements et al., 2011; Dailey et al., 2005; Mu et al., 2012; Teven et al., 2014; Zhang et al., 2014; Zhang, 2009). The MAPK pathway comprises three primary branches, involving phosphorylation of the p38, ERK, and JNK kinases, respectively (Fig. 3a). While JNK has been implicated in ventricular development in the zebrafish embryo (Santos-Ledo et al., 2020) and p38 has a chamber specific role in postnatal hearts (Yokota et al., 2020), little is known about the activation of these kinases in the early embryo (Rose et al., 2010; Wang, 2007; Yokota and Wang, 2016). We therefore investigated whether these individual kinases are activated during gastrulation and whether their activation is affected in *Mesp1* mutant embryos. To this end, we performed western blot analyses for both total and activated forms of the three effector kinases in wild-type and *Mesp1* mutant embryos. We found that all three MAPK pathways are active at both early (OB) and later (LB) stages of gastrulation (Fig. 3b-c, g-h, l-m). For enhanced spatial resolution, we performed immunostaining of the active (phosphorylated) forms of the kinases in the same *Mesp1*− *rtTA/tetO-H2B-GFP* reporter mouse line. At the early bud (EB) stage, p38 and ERK appear to be ubiquitously active throughout the embryo, along the entire proximo-distal axis (Fig. 3d-d,’ i-i’). In contrast, JNK activity is spatially restricted to the posterior-proximal region of the embryo, specifically in the area where posterior SHF progenitors emerge from the PS (Fig. 3n-n’) (Bertrand et al., 2011; Ivanovitch et al., 2021; Lescroart et al., 2018). Our findings thus demonstrate that the p38, ERK and JNK effectors of the MAPK pathway are active in gastrulating embryos and in *Mesp1*+ progenitors.

**Figure 3:**
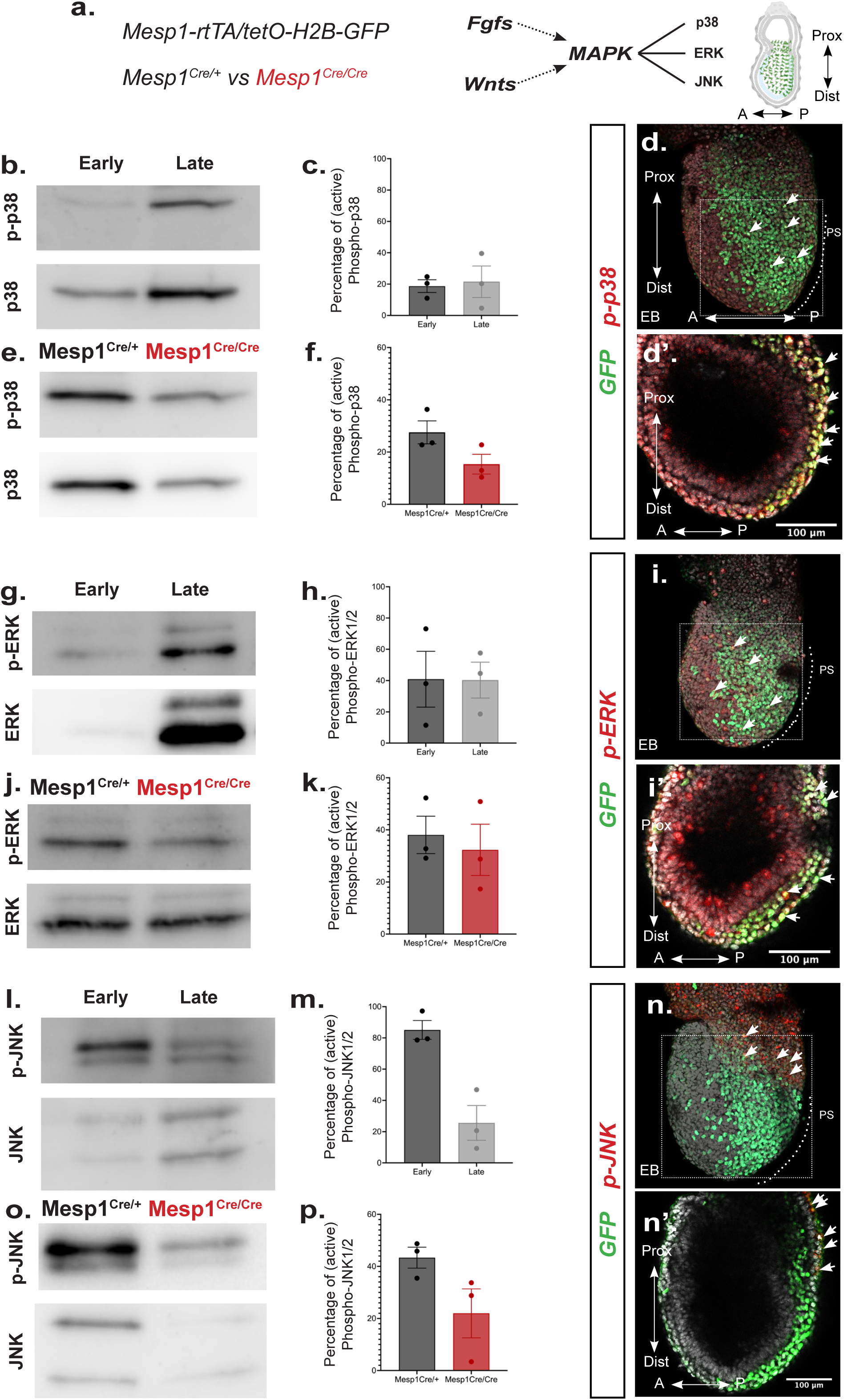
The different branches of the MAPK signalling pathway are active in *Mesp1*+ progenitors during gastrulation and downregulated in *Mesp1* mutants. **a.** Scheme of the experiment. We investigate the activity of different branches of the MAPK pathway in wild-type, *Mesp1^Cre/+^*, *Mesp1^Cre/Cre^* and *Mesp1-rtTA/tetO-H2B-GFP* mouse embryos during gastrulation. **b, e, g, j, l, o.** Western blot analysis of the activity (phosphorylation) of the p38 (~38kDa) (**b** and **e**), ERK1/2 (~42, 44kDa) (**g** and **j**) and JNK1/2 (~46, 54kDa) (**l** and **o**) branches of the MAPK pathway in early (late streak/no bud, E6.75) and late (head fold stages, E7.25) gastrulating embryos (**b**, **g** and **l**) or in *Mesp1* heterozygous (Cre/+) and *Mesp1* mutant (Cre/Cre) embryos (**e**, **j** and **o**). **c, f, h, k, m, p.** Graphs of the percentage of the active (phospho) forms of the MAPK effectors normalized to p38, ERK1/2 or JNK1/2. Results are represented as the mean ± SEM of three biologically independent experiments. Amount of loaded protein was normalized using an antibody against either GAPDH (pERK1/2, ERK1/2, pJNK1/2, JNK1/2) or Actin (pp38, p38). **d, i, n.** Immunostaining analysis of the activity (phosphorylation) of the p38 (**d**), ERK1/2 (**i**) and JNK1/2 (**n**) branches of the MAPK pathway in early bud (EB, E7.25) stage gastrulating embryos. **d’, i’ and n’**. Single optical sections of the embryos in panels d, I and n, magnified as indicated with dotted lines. White arrowheads indicate examples of *Mesp1+* cells (GFP) with MAPK pathway activity. Each embryo is a representative of at least 3 embryos at the defined stage. Scale bars: 100μm. A, anterior; P, posterior; Prox, proximal; Dist, distal; PS, Primitive streak (dotted line). All embryos are represented in the same orientation.

Given the known migration defects of cardiac progenitors in *Mesp1* mutant embryos (Dominguez et al., 2023; Saga et al., 1999) and the observed differences in their signalling environment, we assessed MAPK activity in these embryos by western blot to determine if there was a difference in the activity of the MAPK effectors in *Mesp1* mutant embryos. Due to technical limitations in isolating sufficient numbers of *Mesp1*− expressing cells from mutant embryos, we performed the analysis on the whole embryonic regions from mutant and control embryos. While no significant differences in overall MAPK activity were detected between *Mesp1* mutant embryos and heterozygous controls during gastrulation (Fig. 3e-f, j-k and o-p), stage-dependent differences were observed (Supplementary Fig. 2b-d). p38 and ERK activity appears to be downregulated at earlier stages of gastrulation in *Mesp1* mutant embryos, while JNK activity appears to be affected at a specific stage of gastrulation. Together, these results suggest that MAPK activity is indeed downregulated in *Mesp1* mutant embryos.

### MAPK inhibition affects cardiac progenitor migration

Having established that the MAPK pathway is active during mouse gastrulation, we next aimed to determine its role in the migration of *Mesp1*+ progenitors. To this end, we conducted MAPK inhibition experiments using *ex utero* mouse embryo culture (Fig. 4a). We focused specifically on the p38 and ERK branches of the MAPK pathway, given their widespread activation throughout the embryonic region during gastrulation. Briefly, *Mesp1*− reporter embryos were induced to express GFP at the onset of gastrulation (E6.25). Embryos were dissected shortly thereafter, at early to mid-gastrulation stages (late streak, LS; no bud, OB, Fig. 4a). Following dissection, embryos were first imaged to document their initial morphology (T_0_), and subsequently cultured overnight *ex utero* in the presence of p38 and ERK inhibitors, or DMSO as vehicle control (Supplementary Fig. 4a-c) (Maekawa et al., 2005; Xu et al., 2006)). Embryos were imaged again at 14 to 16 hours post-treatment (T_+16h_), for analysis. Given the established role of MAPK signalling in cell migration, we assessed the extent of *Mesp1*+ progenitor migration, as marked by GFP expression, within the embryonic region. Migration was quantified at two anatomical landmarks: the extra-embryonic/embryonic junction (“Junction”) and the middle of the embryonic region (“Middle”), at both the dissected stages. Measurements were taken at T_+16h_ and normalized to the width of each embryo. Control experiments confirmed that MAPK inhibition did not significantly alter overall size of the embryo, as shown by comparable embryo width (Supplementary Fig. 4d) and length (Supplementary Fig. 4e) between treated and control groups. Importantly, early inhibition (at LS stage) of MAPK signalling resulted in impaired migration of the earliest *Mesp1*+ progenitors, both at the “Junction” and the “Middle” regions (Fig. 4b-f). Notably, simultaneous inhibition of p38 and ERK MAPK branches phenocopied the migration defects observed in *Mesp1* mutant embryos (Supplementary Fig. 4f) (Saga et al., 1999). Interestingly, later inhibition (at OB) selectively impaired the migration of cardiac progenitors emerging in the “Middle” of the embryo (Fig. 4g-k), suggesting a temporally distinct requirement for MAPK signalling during gastrulation. Collectively, these findings indicate that MAPK activity is required for proper *Mesp1*+ progenitor migration, and that compromised MAPK signalling may underlie the migration defects observed in *Mesp1* mutant embryos.

**Figure 4:**
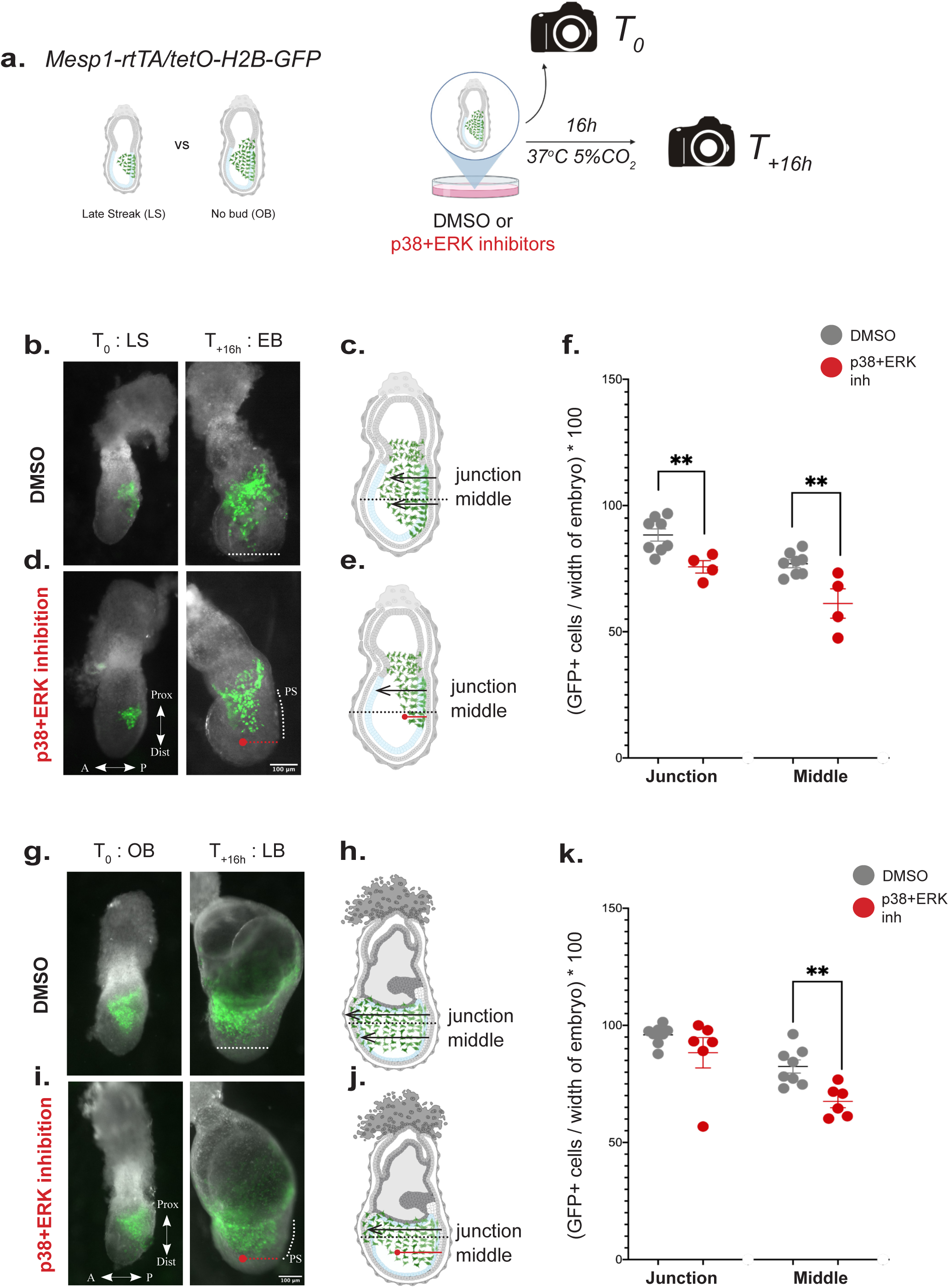
Inhibition of specific MAPK signalling activity dysregulates the migration of *Mesp1*− expressing cells. **a.** Scheme of the experiment. We use *Mesp1* lineage reporter embryos to track *Mesp1* expressing cells at clearly defined embryonic stages of collection (late streak, LS or no bud, OB). Camera icons represent imaging time points at T_0_ and T_+16h_. Gastrulating embryos were cultured *ex utero* in control (DMSO) or MAPK inhibitory (p38+ERK inhibition) conditions. **b-e.** Images of embryos at *T_0_* (late streak stage, LB) and *T_+16h_* and the corresponding representations of later (early bud stage, EB, E6.75) gastrulating embryos cultured *ex utero* in control (DMSO) (**b-c**) or MAPK inhibitory (p38+ERK) conditions (**d-e**). Scale bar: 100μm. **f.** Graph showing the extent of migration of *Mesp1+* (GFP) cells at the embryonic – extra-embryonic junction (junction) or in the middle (middle) of the embryo, normalized to the width of the embryo. Data represented as mean ± SEM. *p* value is calculated using Student’s *t*-test, unpaired, between DMSO and p38+ERK and is indicated above. \*\**p*<0.01. n>3 embryos per conditions. **g-j.** Images of embryos at *T_0_* at the no bud stage (OB) and *T_+16h_* and the corresponding representations of later (LB, late bud, E6.75) gastrulating embryos cultured *ex utero* in control (DMSO) (**g-h**) or MAPK inhibitory (p38+ERK) conditions (**i-j**). Scale bar: 100μm. **f.** Graph showing the extent of migration of *Mesp1+* (GFP) cells at the embryonic – extra-embryonic junction (junction) or in the middle (middle) of the embryo, normalized to the width of the embryo. Data represented as mean ± SEM. *p* value is calculated using Student’s *t*-test, unpaired, between DMSO and p38+ERK and is indicated above. \*\**p*<0.01. n>5 embryos per conditions. A, anterior; P, posterior; Prox, proximal; Dist, distal; PS, Primitive streak (dotted line). All embryos are represented in the same orientation.

### Individual MAPKs play distinct temporal roles in cardiac progenitor development

Given that MAPK inhibition has distinct effects at different stages of early gastrulation, we next investigated whether the requirement for each individual MAPK pathway varies temporally during cardiac progenitor development. This question is particularly relevant, as the emergence of cardiac progenitor subpopulations is temporally segregated. We therefore performed inhibition of each MAPK pathway at systematically and rigorously defined stages of gastrulation (Fig. 5a). For all experiments, embryos were staged according to established morphological criteria (Downs and Davies, 1993). As previously described, *Mesp1*− reporter embryos were generated, with GFP expression induced at either the onset (E6.25) or mid-point (E6.75) of gastrulation. Embryos were subsequently collected at the late streak (LS), no bud (OB), early bud (EB), or late bud (LB) stages. The experimental design was similar to that of the dual inhibition assays described above, but using individual MAPK inhibitors. Embryos collected at LS stage were cultured *ex utero* overnight (T_+16h_) until the EB stage, while those dissected at OB, EB or LB stages were cultured for 6-8 hours (T_+8h_), reaching the EB, LB and early head fold (EHF) stages, respectively. As before, embryos were imaged at both the beginning (T_0_) and end (T_+8h_ or T_+16h_) of the culture period. The impact of each treatment was assessed through quantification of *Mesp1*− GFP+ cell migration and overall embryonic morphology. Control measurements confirmed that, aside from one exception (discussed below), individual MAPK inhibition did not significantly affect embryo size (Supplementary Fig. 4a-c). We first examined the effects of p38 inhibition. *Mesp1+* cardiac progenitors treated with the p38 inhibitor failed to reach the anterior limit observed in control (DMSO-treated) embryos (Fig. 5b-f). The extent of GFP+ cell migration was quantified at the “Middle” of the embryonic region and normalized to embryo width. Notably, inhibition of p38 significantly impaired anterior migration only at early stages of gastrulation (Fig. 5f), indicating that p38 signalling is specifically required during the earliest stages of cardiac progenitor emergence.

**Figure 5:**
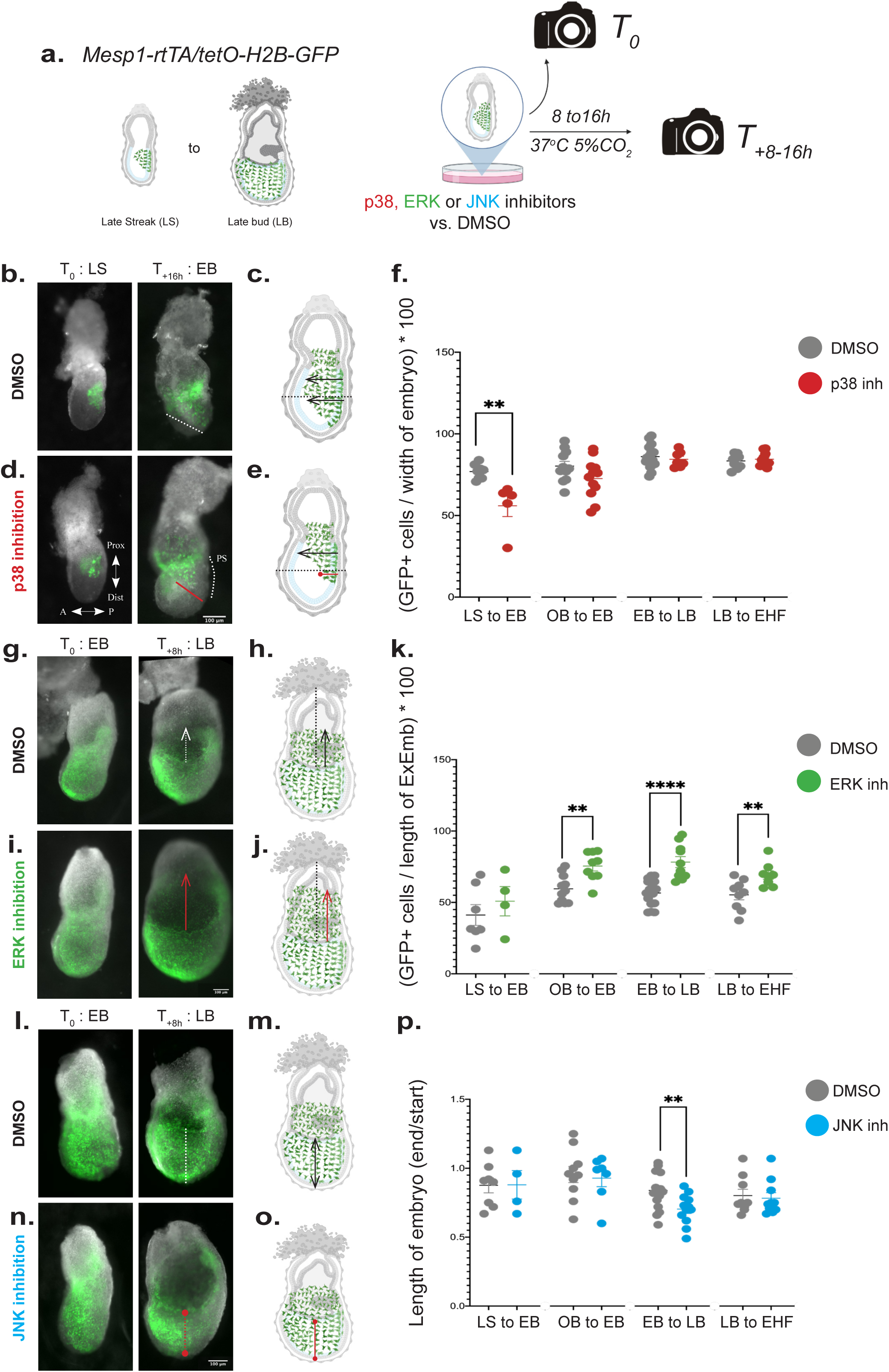
Inhibition of individual MAPK signalling pathways results in distinct phenotypes. **a.** Scheme of the experiment. We use *Mesp1* lineage reporter embryos to track *Mesp1* expressing cells at clearly defined embryonic stages of collection (late streak, LS or early bud, EB). Camera icons represent imaging time points at T_0_ and T_+8-16h_. Gastrulating embryos were cultured *ex utero* in control (DMSO) or MAPK inhibitory (p38 / ERK / JNK inhibition) conditions. **b-e.** Images and corresponding representations of embryos at *T_0_* (late streak stage, LS) and *T_+16h_* and the corresponding representations of later (early bud stage, EB, E6.75) gastrulating embryos cultured *ex utero* in control (DMSO) (**b-c**) or p38 inhibition conditions (**d-e**). Scale bar: 100μm. **f.** Graph showing the extent of migration of *Mesp1+* (GFP) cells in the middle (middle) of the embryo, normalized to the width of the embryo. Data represented as mean ± SEM. *p* value is calculated using Student’s *t*-test, unpaired, between DMSO and p38 inhibition (p38 inh) and is indicated above. \*\**p*<0.01. **g-j.** Images of embryos at *T_0_* (early bud stage, EB) and *T_+8h_* and the corresponding representations of later (late bud stage, LB) gastrulating embryos cultured *ex utero* in control (DMSO) (**g-h**) or ERK inhibition conditions (**i-j**). Scale bar: 100μm. **k.** Graph showing the extent of migration of *Mesp1+* (GFP) cells in the extra-embryonic (ExEmb) region of the embryo, normalized to the length of the extra-embryonic (ExEmb) region of the embryo. Data are represented as mean ± SEM. *p* value is calculated using Student’s *t*-test, unpaired, between DMSO and ERK inhibition (ERK inh) and is indicated above. \*\**p*<0.01. \*\*\*\**p*<0.0001. **l-o.** Images of embryos at *T_0_* (early bud stage, EB) and *T_+8h_* and the corresponding representations of later (late bud stage, LB) gastrulating embryos cultured *ex utero* in control (DMSO) (**l-m**) or JNK inhibition conditions (**n-o**). Scale bar: 100μm. **p.** Graph showing the change in the length of embryonic region at *T_+8h_* of the culture period, normalized to the length of the embryonic region at *T_0_*. Data represented as mean ± SEM. *p* value is calculated using Student’s *t*-test, unpaired, between DMSO and JNK inhibition (JNK inh) and is indicated above. \*\**p*<0.01. A, anterior; P, posterior; Prox, proximal; Dist, distal; PS, Primitive streak (dotted line). All embryos in this figure are visualized in the same orientation.

We then assessed the role of the ERK pathway. While the antero-lateral migration of *Mesp1*+ cardiac progenitors was not significantly altered in ERK-inhibited embryos compared to controls, we observed an extension of the GFP+ domain further into the extra-embryonic region (Fig 5g-j). This effect was quantified from the extra-embryonic/embryonic junction and normalized to the length of the extra-embryonic region. Although a trend was seen with early inhibition, a significant phenotype became apparent only from mid-gastrulation stages (OB) onwards (Fig. 5k). Western blot analysis for phosphor-histone-H3 (pHH3) revealed no significant change in proliferation between treated and control embryos (Supplementary Fig. 4d-e). These findings indicate that ERK signalling regulates *Mesp1*+ progenitor migration in the extra-embryonic region, particularly from mid-gastrulation stages onward.

Finally, we investigated the role of JNK signalling. Inhibition of JNK resulted in embryos with shorter embryonic region at a specific stage of gastrulation (Supplementary Fig. 4b-c). Embryonic length was measured at both T_0_ and T_n_ (T_+8h_ or T_+16h_), and normalized to the total embryo length (embryonic + extra-embryonic) (Fig. 5l-o). A ratio of “End” to “Start” embryonic length was used to estimate proximo-distal axial elongation. A significant effect was observed only at the EB stage, where JNK inhibition impaired axial growth (Fig. 5p). At this same stage, *Mesp1*+ progenitors at the “Middle” of the embryo also displayed reduced antero-lateral migration (Supplementary Fig. 4h-i), paralleling the stage-specific reduction in JNK activity observed in *Mesp1* mutant embryos (Supplementary Fig 2d). These results suggest that JNK activity is required at a specific time point during gastrulation to regulate both axial growth and *Mesp1*+ progenitor behaviour. Taken together, these findings demonstrate a clear temporal heterogeneity in the requirement for MAPK pathway components during cardiac progenitor development: p38 signalling is critical at the onset of gastrulation, JNK is required at later stages, and ERK appears to play a role throughout gastrulation.

## Discussion

Our results provide evidence that the previously reported heterogeneity of signalling ligands among cardiac progenitor populations during gastrulation (Lescroart et al., 2018, 2014) extends to the downstream intracellular signalling effectors. Inhibition of these effectors during gastrulation phenocopies the well-characterized migration defects observed in *Mesp1* mutant embryos (Dominguez et al., 2023; Saga et al., 1999). Notably, individual inhibition of each downstream MAPK effector reveals distinct temporal requirements for the normal migration of *Mesp1*+ progenitors. This temporal specificity aligns with the sequential emergence of cardiac progenitor populations, each contributing to specific cardiac regions (Fig. 6).

**Figure 6:**
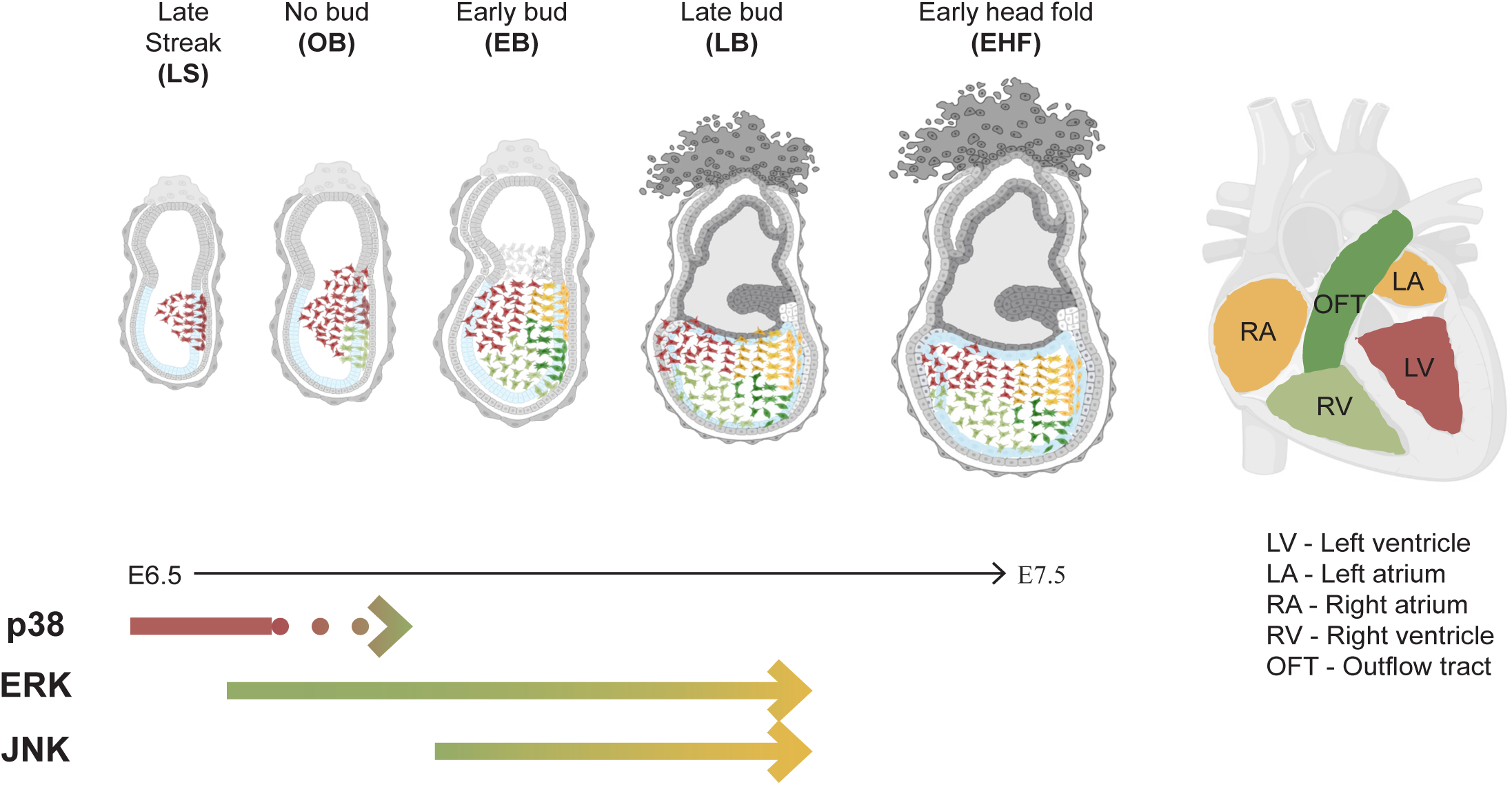
MAPK effectors are required for cardiac progenitor migration in distinct temporal windows during gastrulation. Scheme of the temporal requirement of MAPK effectors during *Mesp1+* cardiac progenitor emergence and migration.

We first provide evidence that distinct cardiac subpopulations are exposed to a heterogenous signalling microenvironment. Our analysis of candidate intercellular signalling ligand gene expression (Fig. 1), together with previous studies (Arnold and Robertson, 2009; Chen et al., 2019; Du et al., 2010; Kimura et al., 2001; Kinder et al., 1999; Niswander and Martin, 1992; Nowotschin et al., 2012; Sun et al., 1999; Tremblay et al., 2001; Yamaguchi et al., 1999), suggests that this heterogeneity may reflect the dynamic and continuously evolving nature of the signalling microenvironment during gastrulation. Single cell RNA-sequencing data (Lescroart et al., 2018) further indicates that *Mesp1* mutant embryos display an altered signalling niche during gastrulation. Our results confirm that even the subset of pro-migratory ligand encoding genes we tested (from the FGF and WNT pathways) exhibits such differences, suggesting that changes in ligand availability may contribute to the migration defect observed in *Mesp1* mutants embryos (Dominguez et al., 2023; Saga et al., 1999).

Despite the heterogeneity in ligand expression, relatively few receptors appear to mediate signal transduction from multiple ligands during gastrulation. One key receptor is FGFR1, whose inactivation leads to severe migration defects in mouse embryos with early emerging cells accumulating in the PS (Yamaguchi et al., 1994). The cellular responses triggered by the ligand-receptor interactions are ultimately executed by intracellular signalling effectors. These effectors are of particular interest, as they not only relay signals from upstream components but can also act as integration nodes for multiple signalling pathways, enabling coordinated cellular responses (Bikkavilli and Malbon, 2009; Clements et al., 2011; Dailey et al., 2005; Mu et al., 2012; Qin et al., 2024; Zhang et al., 2014; Zhang, 2009, p. 200). MAP kinases are well-established intracellular signalling effectors known to regulate cell fate decisions (Peterson et al., 2022), proliferation (Zhang and Liu, 2002) and migration (Huang et al., 2004), all key behaviours of cardiac progenitors during gastrulation. Although the signal transduction mechanisms (e.g., phosphorylation of subsequent kinases) are broadly conserved across the MAPK branches, the specificity of the cellular response is determined by the precise combination of ligand, receptor, adaptor, and scaffold proteins. Our data confirms that MAP kinases are active in cardiac progenitors during gastrulation and that their activity is reduced in *Mesp1* mutant embryos in a stage-specific manner. p38 and ERK activity appears diminished at early stages of gastrulation, while JNK activity is affected at a specific stage. However, due to technical limitations in isolating cardiac progenitors from each individual mutant embryo, MAPK activity was measured in the entire embryonic region using western blot analysis. This approach may have masked more subtle or lineage specific differences in kinase activation between *Mesp1* mutant embryos and controls. To overcome this limitation, we employed pharmacological inhibition. Notably, simultaneous inhibition of two MAPK effectors recapitulates the migration defect observed in *Mesp1* mutants (Saga et al., 1999). This is particularly striking given the heterogeneous expression of upstream MAPK ligands in control embryos (Fig 1) and their dysregulation in the mutants (Fig 2, (Lescroart et al., 2018). Together, these findings suggest that impaired MAPK signalling may be a contributing mechanism underlying the migration defects observed in *Mesp1* mutant embryos. Interestingly, an interplay between glucose metabolism, FGF ligands, and ERK signalling has been established in controlling cell migration during mouse gastrulation (Cao et al., 2024). However, a potential link between glucose metabolism and the behaviour or fate of the distinct cardiac subpopulations remains to be clarified.

Although the various branches of the MAPK pathway are clearly active during gastrulation, our stage-resolved analyses unexpectedly reveal that their functional requirement is restricted to distinct temporal windows (Fig. 6). This temporal specificity may reflect the heterogeneity of upstream intercellular signalling ligands that converge on each MAPK effector. Our data indicate that p38 and ERK activity is required early, during the emergence of the FHF/JCF, while ERK and JNK are required later, coinciding with the specification and migration of the SHF. This model is supported by evidence from other genetic studies. In mutants for a p38 interacting/activating protein, early mesoderm – including FHF/JCF progenitors – accumulates at the PS due to a failure to downregulate E-cadherin, impairing migration (Zohn et al., 2006), a phenotype similar to that observed in *Fgfr1* and *Fgf8* mutant embryos, as well as embryos mutant for the WNT co-receptor *Lrp5/6* (Deng et al., 1994; Kelly et al., 2004; Sun et al., 1999; Yamaguchi et al., 1994). Conversely, in *Rac1* mutant embryos, where Rac1 acts downstream of JNK, cardiac mesoderm is specified (as shown by *Nkx2.5* expression) but fails to complete its anterior migration, resulting in cardiac bifida (Migeotte et al., 2011). This phenotype closely resembles that of milder *Mesp1* mutants that also display cardiac bifida (Saga et al., 1999), where later-emerging mesodermal populations appear affected. Whether MAPK effectors serve as true integrative hubs that convert multiple upstream signals into context-specific cellular responses remains an open question. Testing this hypothesis would require gain- or loss-of-function manipulations of individual MAPK branches specifically in defined cardiac progenitor subpopulations. However, such experiments remain technically challenging due to the need for precise spatial and temporal control, as well as the difficulty of modulating kinase activity, which typically depends on tightly regulated phosphorylation events.

The effects of MAP kinase inhibition reported here are robust and reproducible, having been observed across multiple independent experiments, litters and gastrulation stages. As is standard in the field, inhibitor treatments were applied globally in *ex utero* embryo cultures. While this approach does not allow us to distinguish between cell-autonomous and non-cell-autonomous effects, the data nonetheless reveals clearly defined temporal windows during which a specific MAPK effector is required. These windows align with the sequential emergence of distinct cardiac progenitor subpopulations.

Collectively, our results provide the first evidence that heterogeneity of the signalling niche at the time of *Mesp1+* cardiac progenitor emergence may correlate with the temporal heterogeneity in the requirement for specific downstream intracellular signalling effectors. These effectors likely act as integrators of multiple upstream signals, coordinating cellular responses such as fate specification and migration – either by directly influencing cell fate-determining networks or by modulating the spatial positioning of progenitors within defined signalling environments. Further studies will be required to disentangle these possibilities, combining cell fate markers with the pathway inhibition, and employing live imaging to determine whether *Mesp1+* progenitors adopt distinct migratory trajectories in MAP kinase inhibited embryos. A comprehensive understanding of how cardiac chamber progenitor fates are specified is not only critical for elucidating the principles of heart development and etiology of CHDs, but is also essential for improving protocols to generate specific cardiac cell types from pluripotent stem cells.

## Acknowledgements

We thank C. Blanpain for providing the *Mesp1-rtTA/tetO-H2B-GFP* mouse line. We thank the MMG*IP Photonic imaging platform and the Animal Phenotyping Core platform (MMG). We thank Robert Kelly, Christopher De Bono and Stéphane Zaffran for careful reading of the manuscript. F.L.’s laboratory was supported by the INSERM ATIP-Avenir program. N. N. was funded by the Fondation pour la Recherche Médicale (FRM) [SPF202005012035]. S. E.I. was founded by a fellowship from the French ministry of research.

## Materials and methods

### Mouse lines

CD1/Swiss mice were used as wildtype. *Mesp1-rtTA/TetO-H2B-GFP* (Lescroart et al. 2014) and *Mesp1^Cre/+^* (Saga et al. 1999) mice have been previously described and were obtained from the laboratory of Cédric Blanpain. To generate *Mesp1^Cre/Cre^; Mesp1-rtTA/TetO-H2B-GFP* embryos, *Mesp1^Cre/+^* mice were bred into the *Mesp1-rtTA/TetO-H2B-GFP* background. The colonies were maintained in a certified animal facility (agreement #C 13 013 08) in accordance with European guidelines. The local ethical committee (CEBEA) approved the experiments and they comply with the relevant ethical regulations concerning animal research (Ministère de l’Education Nationale, de l’Enseignement Supérieur et de la Recherche; Authorization N 32-08102012).

### Embryo dissections

Gastrulation stage embryos were dissected in the evening on the 6^th^ day after observation of a vaginal plug (E0.5) to obtain E6.75. Embryos were dissected on the morning of the 7^th^ day for E7.25 and early in the afternoon of the 7^th^ day for E7.5/E7.75. Despite these dissection time points, the embryos were precisely staged at the time of dissection, using the morphological landmarks described in Downs and Davies 1993 (Downs and Davis 1993), to describe the stage as mid-streak (MS), late streak (LS), no bud (OB), early bud (EB), late bud (LB), early head fold (EHF) or late head fold (LHF). Reporter gene expression was induced by 40mg/kg intra-peritoneal injection of doxycycline (Dox) at E6.25 for dissection at E6.75 or at E6.75 for dissection at E7.25, E7.5 or E7.75. Embryos were carefully staged at dissection and, whenever possible, staging was reconfirmed at the time of analysis. All results are based on a minimum of three embryos per condition and compared to stage- and litter-matched controls.

### *Ex utero* embryo cultures

Embryos were dissected in dissection medium containing DMEM/F12 (ThermoFisher Scientific, #11320074) with 10% heat inactivated fetal bovine serum (Pansera, #P30-2602), supplemented with 1X penicillin-streptomycin (Gibco, #15070-063). An Axiozoom v16 microscope was used to select GFP expressing embryos. These embryos were then stage-matched and washed in embryo culture medium containing DMEM/F12 (ThermoFisher Scientific, #21041033) supplemented with 50% heat inactivated rat serum (Janvier) before being transferred into individual wells of a 15-well angiogenesis slide (ibidi, #81506) containing the same embryo culture medium supplemented with vehicle (DMSO, Invitrogen, D12345) or one or a combination of the MAPK inhibitors (20μM p38 inhibitor – SB203580, Calbiochem #559389], 50μM ERK inhibitor – U0126, CST #9903S, 25μM JNK inhibitor – JNK inhibitor II, Calbiochem #420119). The embryos dissected at E6.75 were cultured overnight (~14-16 hours), while those dissected at E7.25 were cultured for 6-8 hours at 37°C with 5% CO_2_ as a static culture. All embryo measurements were performed in Fiji/ImageJ.

### Whole mount RNAscope

Embryos were dissected in PBS and rinsed once with PBS before being incubated in 4% PFA at 4°C for 36-48 hours. Embryos were dehydrated in a progressive methanol series and stored in 100% methanol. For further processing, the embryos were rehydrated in an inverse methanol series. The embryos were subsequently processed using an adapted version of the protocol of the RNAscope Multiplex Fluorescent v2 kit (ACD-Bio #323110). *Mm-Fgf3-C1* (503101-C1), *mm-Fgf4-C1* (514311-C1), *mm-Fgf8-C2* (313411-C2), *mm-Wnt3-C1* (312241-C1), *mm-Wnt3a-C2* (405041-C2), *mm-Wnt5a-C1* (316791-C1), *mm-Hand1-C3* (429651-C3), *mm-Foxc2-C1* (412851-C1), *mm-Hoxb1-C3* (541861-C3) and *mm-EGFP-C4* (400281-C4) were the probes used. Embryos were imaged in PBS using the LSM800 (Zeiss) confocal microscope or on a Lightsheet 7 (Zeiss). Individual embryos were then collected and only the embryonic region was used to genotype the embryos using the Phire Tissue Direct PCR Master Mix (ThermoFisher Scientific, F170S).

### Western blot

The embryonic region of stage-matched wildtype embryos was pooled and lysed using RIPA buffer (ThermoFisher Scientific, #89900) containing a protease and phosphatase inhibitor cocktail (ThermoFisher Scientific, #78442) and sonicated using the Diagenode Bioruptor (UCD-200). For mutant embryo litters, individually labelled extra-embryonic regions were collected for genotyping while embryonic regions were collected in correspondingly labelled tubes for lysis and sonication. After genotyping, litter and stage-matched embryos of identical genotypes were pooled. Protein concentration was measured using the Pierce protein assay reagent (#22660). 900ng-2ug of sample was loaded on a precast gel (Novex, #XP04202BOX) and separated according to size. The protein was transferred to a nitrocellulose membrane and blocked for 1 hour in 5% BSA (Sigma Aldrich, A7906) in 0.1% Tween-20 containing TBS (TBST). Specific primary antibodies diluted in the same blocking solution were incubated with the membrane overnight at 4°C and washed six times with TBST. The membrane was then incubated with HRP-coupled secondary antibody for 1 hour before being washed six times with TBST. Revelation was done using an ECL kit (BioRad Clarity, #1705060) in a BioRad Chemidoc MP Imaging System. Primary antibodies used were anti-phospho-p38 (Cell Signalling Technologies, # 9211, 1:1000), anti-p38 (Cell Signalling Technologies, # 8690, 1:1000), anti-phospho-p44/42 MAPK (ERK1/2) (Cell Signalling Technologies, # 4376, 1:1000), anti-p44/42 MAPK (ERK1/2) (Cell Signalling Technologies, # 4696, 1:2000), anti-phospho-SAPK/JNK (Cell Signalling Technologies, # 9255, 1:500), anti-SAPK/JNK (Cell Signalling Technologies, #9252, 1:500), anti-phospho-cJun (Cell Signalling Technologies, #9164, 1:500) and anti-phospho-HH3 (Sigma Aldrich, #06-570, 1:500) and secondary antibodies used were anti-mouse HRP (ThermoFisher Scientific, #A16011, 1:5000) and anti-rabbit HRP (ThermoFisher Scientific, #A16096, 1:7500). Quantity of loaded protein was normalized using an anti-Actin (Sigma Aldrich, MAB1501R, 1:500) or anti-GAPDH (Sigma Aldrich, MAB374, 1:5000) antibody. Fiji/ImageJ was used to quantify the ECL signals and all graphs represent three independent biological experiments.

### Immunofluorescence assay

Embryos were dissected in PBS and after a rinse of PBS were fixed using 4% PFA at 4*C for 45 minutes. After a wash in PBS with 0.1% Tween-20, the embryos were permeabilized with 3 washes of PBS containing 0.3% Triton-X (PTx). Embryos were blocked for 24 hours at 4°C in PTx containing 2% BSA. Primary antibody incubation was performed at 4°C for 60-72 hours in PTx containing 2% BSA. Primary antibodies used were anti-chicken GFP (Thermofisher, A10262, 1:200) with anti-phospho-p38 (Cell Signalling Technologies, #9211, 1:100), anti-phospho-p44/42 MAPK (ERK1/2) (Cell Signalling Technologies, #4376, 1:400) or anti-phospho-SAPK/JNK (Cell Signalling Technologies, #9255, 1:100). After washing for 5-6 hours in PBS containing 0.15% Tween-20 and 2% BSA (PTw+BSA) the embryos were incubated with secondary antibodies anti-rabbit 647 (ThermoFisher Scientific, #A32733, 1:500) or anti-mouse 647 (ThermoFisher Scientific, #A-21235, 1:500), anti-chicken 488 (ThermoFisher Scientific, #A78948, 1:500) and DAPI (ThermoFisher Scientific, #62248, 1:1000). Embryos were subsequently washed for 5-6 hours in PTw+BSA and imaged in PBS at an LSM800 (Zeiss) confocal microscope.

