## Supplemental Figures for "Temporal requirements of MAPK effectors reflect signalling microenvironment heterogeneity during *Mesp1*+ cardiac progenitor emergence and migration"

### Supplementary Figure 1

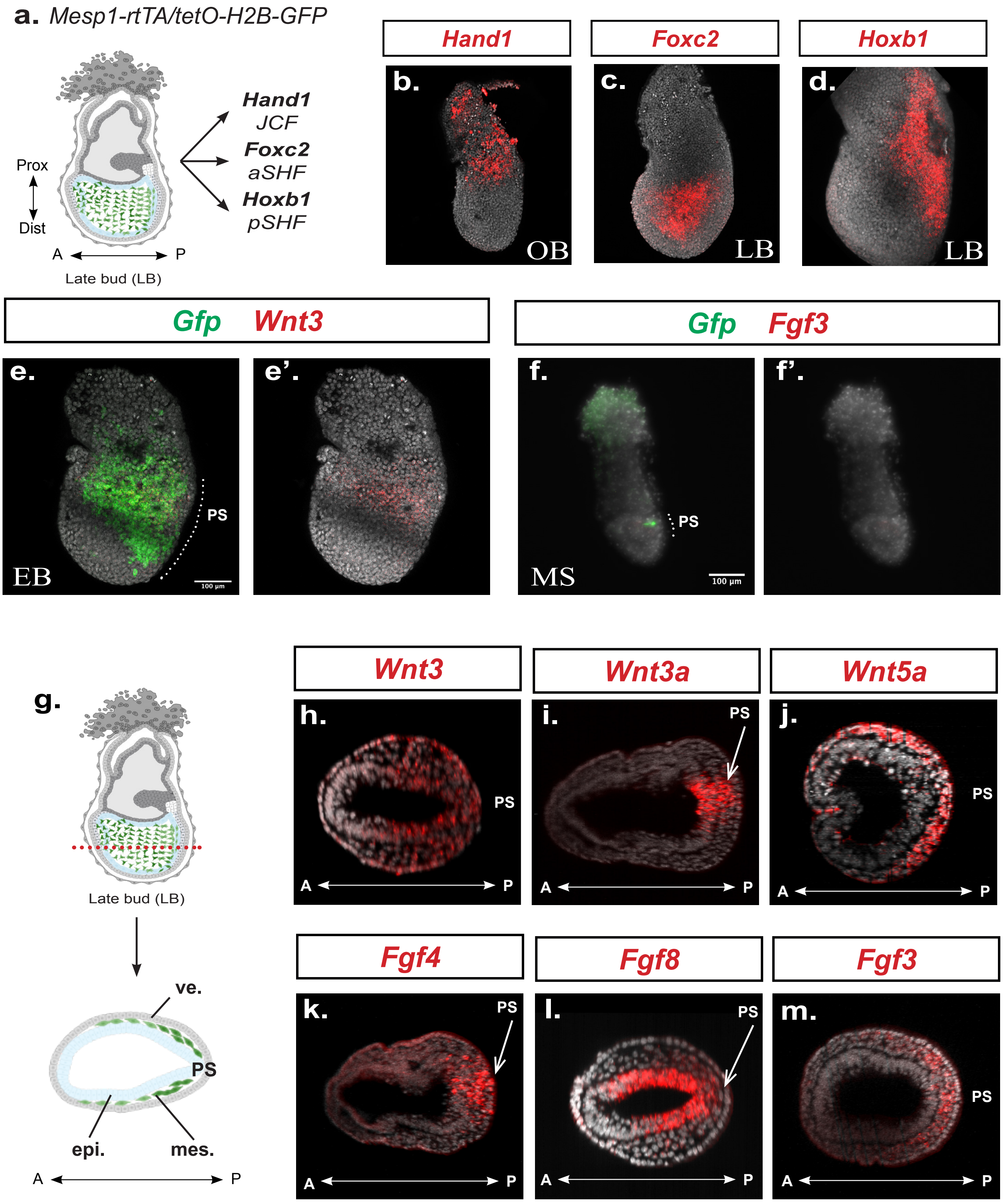

**Supplementary Figure S1: The signalling environment experienced by *Mesp1*+ progenitors evolves during gastrulation.**

**a.** Scheme of the experiment. **b-d.** Representative maximum intensity projection of embryos after RNAscope with *Hand1* (**b**), *Foxc2* (**c**) or *Hoxb1* (**d**) probes. **e-f.** Representative maximum intensity projection of embryos after RNAscope with *Gfp* and *Wnt3* (**e**) or *Fgf3* (**f**) probes. **e'-f'** represent the same images with only the expression pattern of *Wnt3* or *Fgf3*. Each embryo is a representative of at least 3 embryos. Scale bar: 100μm. **g.** Scheme of a transverse section of an embryo at E7.5. ve,, visceral endoderm; epi., epiblast; mes., mesoderm; PS, Primitive streak. **h-m.** Single transverse optical sections demonstrating the expression of the ligands *Wnt3* (**h**), *Wnt3a* (**i**), *Wnt5a* (**j**), *Fgf8* (**k**), *Fgf4* (**l**) and *Fgf3* (**i**) in each germ layer of the embryo at gastrulation. A, anterior; P, posterior; PS, Primitive streak.

### Supplementary Figure 2

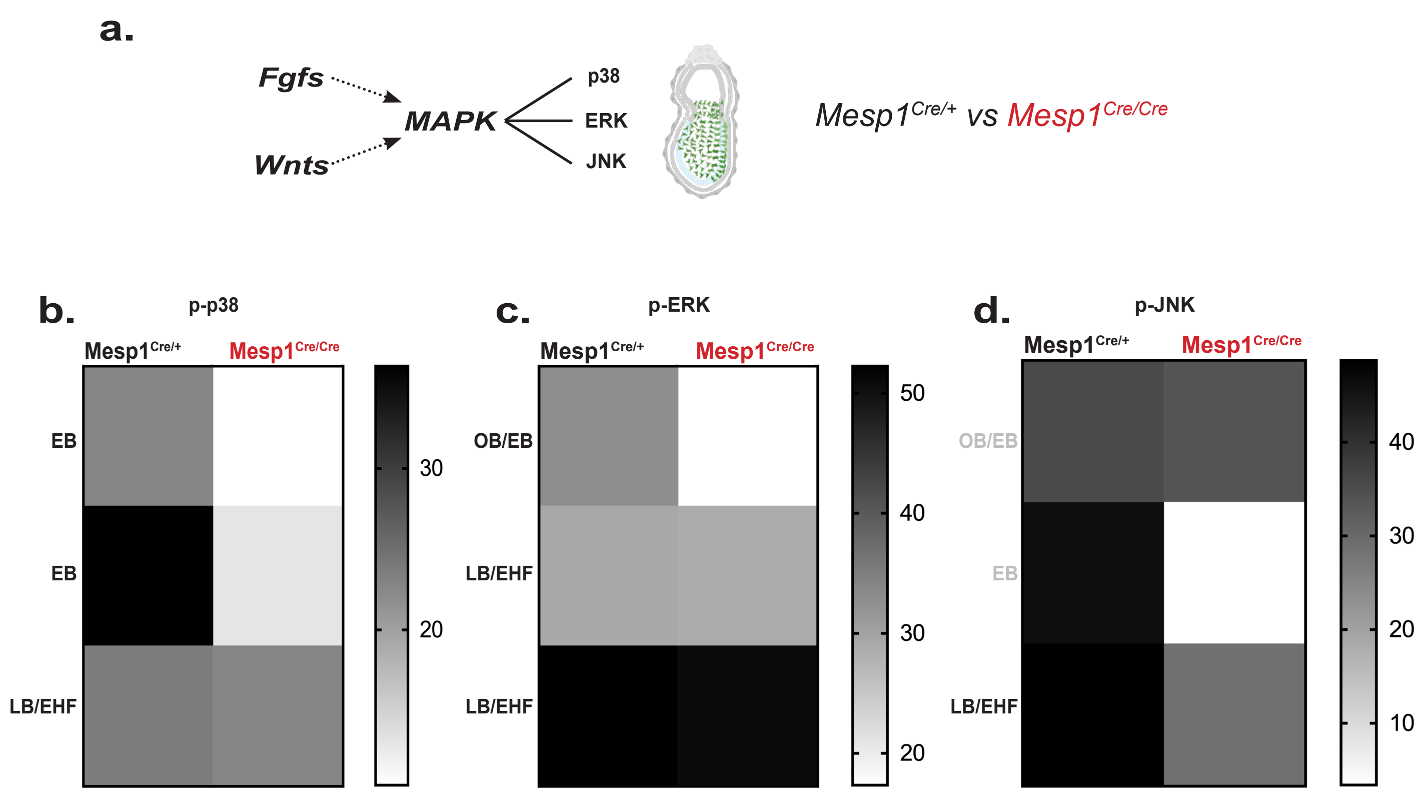

**Supplementary Figure S2: Downregulation of MAPK activity in *Mesp1* mutants appears to be correlated with developmental stage.**

**a.** Scheme of the experiment. **b-d.** Heatmap representation of the activity (phosphorylation) of the p38 **(b)**, ERK1/2 **(c)** and JNK1/2 **(d)** branches of the MAPK pathway in *Mesp1* heterozygous (*Cre/*+) and *Mesp1* mutant (*Cre/Cre*) embryos, segregated by stage. These are individual data points of the data presented in Fig. 3f, k and p.

**Supplementary Figure 3**

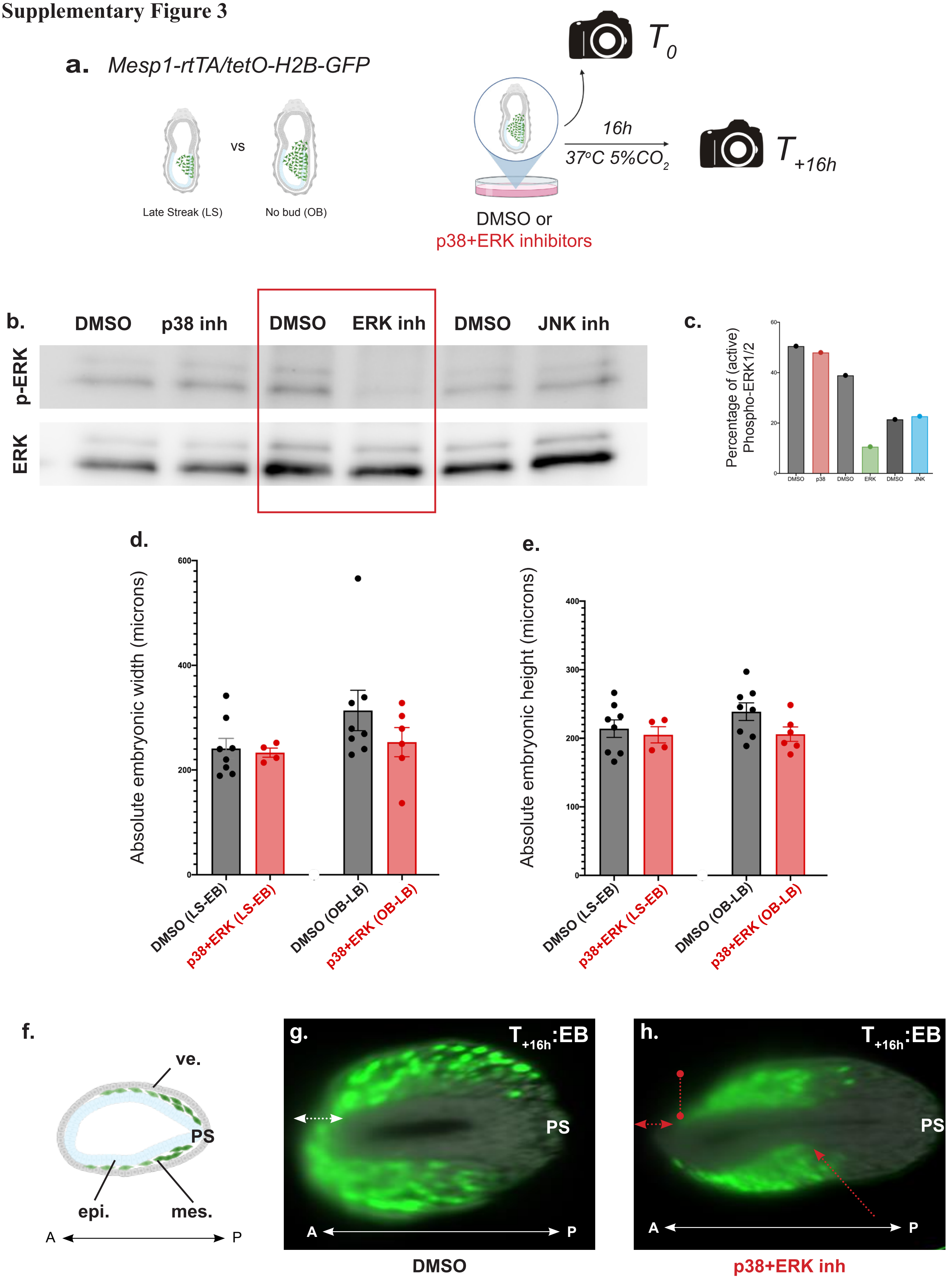

**Supplementary Figure S3: MAPK inhibition is specific to the pathway branch and does not seem to affect overall morphology of the embryos.**

**a.** Scheme of the experiment. **b.** Western blot analysis of the activity (phosphorylation) of the ERK1/2 branch of the MAPK pathway in control (DMSO) or p38, EKR1/2 or JNK1/2 inhibited embryos. Amount of loaded protein was normalized using an antibody against GAPDH. **c.** Graph of the western blot analysis in (**b**) of the active form of the ERK1/2 branch. **d-e.** Graph depicting the embryo morphology analysed by measuring the width (**d**) and length (**e**) of the embryonic region in control (DMSO) or MAPK inhibitory (p38+ERK) conditions. Data is represented as mean ± SEM. *p* value is calculated using Student's *t*-test, unpaired, between DMSO and p38+ERK, but the *p* value is greater than 0.05 and is not indicated on the graph. **f.** Scheme of a transverse section of an embryo at E7.5. ve,, visceral endoderm; epi., epiblast; mes., mesoderm; PS, Primitive streak. **g-h.** Single transverse optical sections demonstrating the extent of anterior migration of *Mesp1*+ (GFP) cells in control (DMSO) or MAPK inhibitory (p38+ERK) conditions at $T_{+16h}$ of *ex utero* culture. **g**. White ←→ line represents presence of *Mesp1*-GFP+ cells anterior to the anterior epiblast. **h**, Red ←→ line (horizontal) represents absence of *Mesp1*-GFP+ cells anterior to the anterior epiblast and vertical red line indicates the anterior limit of the *Mesp1*-GFP+ cells. The red arrow indicates accumulation of *Mesp1*-GFP+ cells towards the posterior. A, anterior; P, posterior; PS, Primitive streak.

**Supplementary Figure 4**

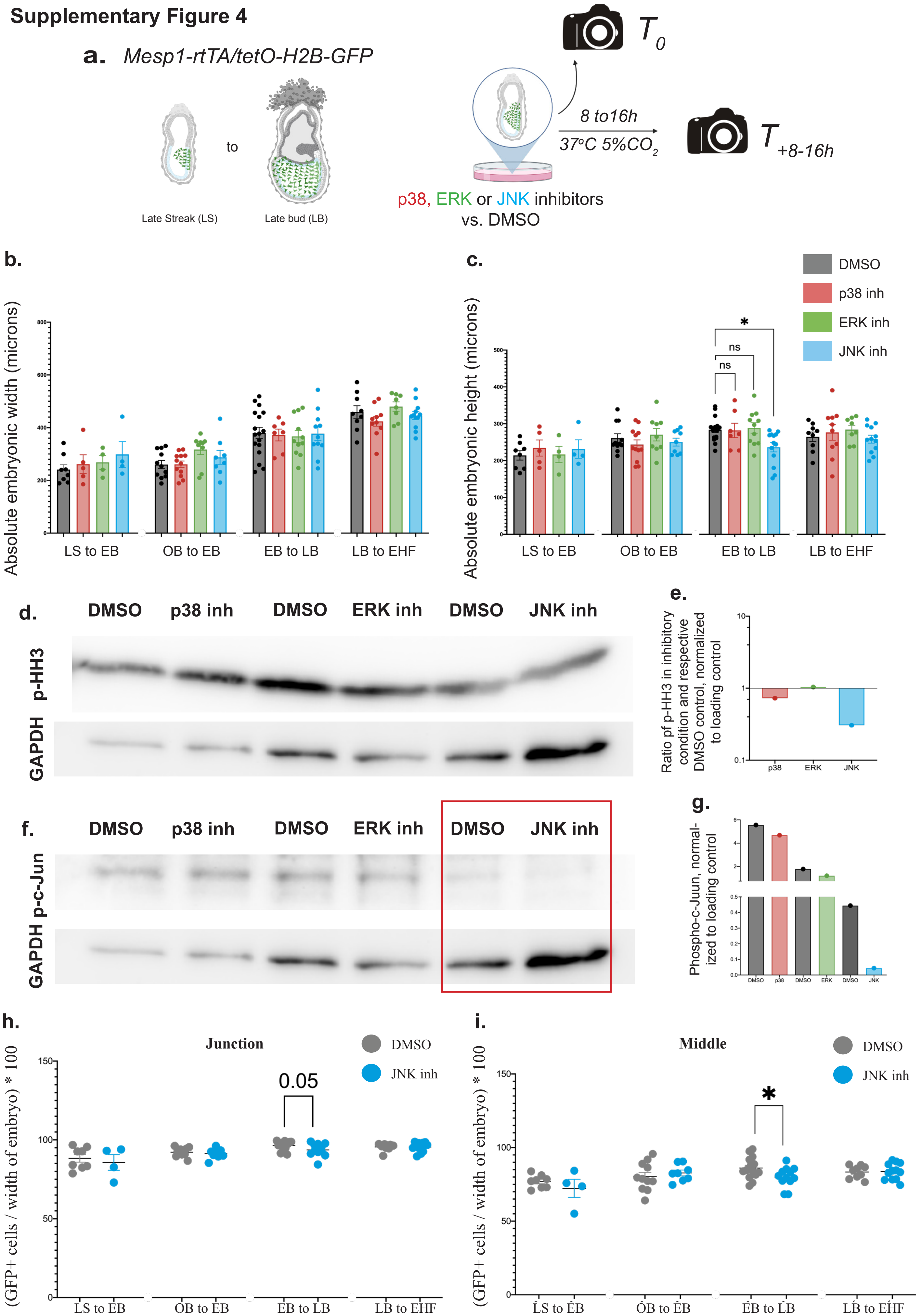

**Figure S4: MAPK inhibition and its consequent effects appear to be specific to the pathway branch**

**a.** Scheme of the experiment. **b-c.** Graph depicting embryo morphology by measuring the width (**b**) and length (**c**) of the embryonic region in control (DMSO) or MAPK inhibitory (p38, ERK1/2 or JNK1/2 inhibition) conditions. Data is represented as mean ± SEM. *P* value is calculated using one-way ANOVA followed by a Dunnett's post-hoc test comparing each inhibitory condition to DMSO. *P* value for (**b**) is greater than 0.05 and is not indicated on the graph. *P* value for (**c**) is less than 0.05 and is not indicated on the graph, but the specific comparison with $p<0.05$ is indicated with *. ns, not significant. **d.** Western blot analysis of the proliferation (phosphorylation of Histone H3) in control (DMSO) or p38, EKR1/2 or JNK1/2 inhibited embryos. Amount of loaded protein was normalized using an antibody against GAPDH. **e.** Graph of the western blot analysis in (**d**) of proliferation, normalized to the control (DMSO) value. **f.** Western blot analysis of the activity of the JNK1/2 branch of the MAPK pathway (phosphorylation of c-Jun, a JNK pathway downstream component) in control (DMSO) or p38, EKR1/2 or JNK1/2 inhibited embryos. Amount of loaded protein was normalized using an antibody against GAPDH. Both p-c-Jun (48kDa) and p-HH3 (17kDa) were probed on the same blot so the loading control (GAPDH, 36kDa) is the same. **g**. Graph of the western blot analysis in (**f**) of the active form of c-Jun, the downstream component of the JNK1/2 branch. **h-i.** Graph showing the extent of anterior migration of Mesp1+ (GFP) cells at the embryonic – extra-embryonic junction (junction, **h**) or in the middle (middle, **i**) of the embryo, normalized to the width of the embryo. Data represented as mean ± SEM. *p* value is calculated using Student's *t*-test, unpaired, between DMSO and JNK inhibition and is indicated above. *$p<0.05$.
